# Intra-host viral populations of SARS-CoV-2 in immunosuppressed patients with hematologic cancers

**DOI:** 10.1101/2022.10.19.512884

**Authors:** Dominique Fournelle, Fatima Mostefai, Elsa Brunet-Ratnasingham, Raphael Poujol, Jean-Christophe Grenier, José Héctor Gálvez, Amélie Pagliuzza, Inès Levade, Sandrine Moreira, Simon Grandjean Lapierre, Nicolas Chomont, Daniel E. Kaufmann, Morgan Craig, Julie G. Hussin

## Abstract

Throughout the SARS-CoV-2 pandemic, several variants of concern (VOC) have been identified, many of which share recurrent mutations in the spike protein’s receptor binding domain (RBD). This region coincides with known epitopes and can therefore have an impact on immune escape. Protracted infections in immunosuppressed patients have been hypothesized to lead to an enrichment of such mutations and therefore drive evolution towards VOCs. Here, we show that immunosuppressed patients with hematologic cancers develop distinct populations with immune escape mutations throughout the course of their infection. Notably, by investigating the co-occurrence of substitutions on individual sequencing reads in the RBD, we found quasispecies harboring mutations that confer resistance to known monoclonal antibodies (mAbs) such as S:E484K and S:E484A. Furthermore, we provide the first evidence for a viral reservoir based on intra-host phylogenetics. Our results on viral reservoirs can shed light on protracted infections interspersed with periods where the virus is undetectable as well as an alternative explanation for some long-COVID cases. Our findings also highlight that protracted infections should be treated with combination therapies rather than by a single mAbs to clear pre-existing resistant mutations.

## Introduction

Several SARS-CoV-2 VOCs have convergent mutations in the spike protein’s RBD that coincide with known epitopes.^1^ Mutations in this genomic region affect the ability of the spike (S) to enter the cell via the ACE2 receptor and have been linked with higher transmission rates and/or immune escape.^2,3^

While in most cases, SARS-CoV-2 infections are cleared within a few days, key mutations develop *de novo* in long lasting infections in patients with immunosuppressive conditions. These infections can last for several months, and their viral mutation rate is higher than in shorter infections in immunocompetent patients.^4^ For this reason, it is suspected that protracted infections are one of the drivers of SARS-CoV-2’s genomic evolution and a source of immune escape variants.^5^ One such example is S:E484K that was found in former VOCs Beta and Gamma.^6^ This mutation has been shown to give the virus immune escape properties such as resistance to anti-viral monoclonal antibodies (mAbs) and convalescent sera as well as reinfection.^7^ Resistance to these treatments has become a growing concern during the past year as an increasing number of Omicron sub-lineages were found to be resistant to a variety of mAbs ^8,9^

Despite the potential importance of these cases, few longitudinal datasets of sequences collected from immunosuppressed individuals at different time points during their infections are available. These datasets can give us insight into evolutionary events that are not observed in acute infections, such as an instance of recombination between two viral strains^10^ or the presence of distinct viral populations with immune escaping mutations in a single sample.^11^

Here, we describe the genetic events that arose in two patients with hematologic cancers that were infected by SARS-CoV-2 for several months. Samples from the first patient (Q1) were collected by the Public Health Laboratory of Québec (LSPQ), in Canada. Viral sequences from the second patient (K1) were generated by Lee et al.^12^ from samples collected in Korea. Through phylogenetic and intra-host single nucleotide variant (iSNV) analysis, we show evidence for a mutational pattern suggestive of a viral reservoir as well as for several viral populations containing immune escape mutations in the spike’s RBD.

## Results

### Description of patients

An immunosuppressed 73-year-old woman (Q1) with non-Hodgkin lymphoma first tested positive for SARS-CoV-2 (PANGO lineage B.1.160) on 08/01/2021 (Day 1, D1). She had undergone several courses of anti-CD20 (rituximab) and chemotherapy in the months preceding her COVID diagnosis. She was vaccinated with the Pfizer vaccine on 25/02/2021. The patient tested positive again on 28/04/2021 (D111). The full timeline of her infection is shown in Figure 1a. Because the sample sequenced on D111 had S:E484K, it was first assumed that this sample and all subsequent timepoints were from a reinfection. However, phylogenetic analysis of all time points shows that all samples came from the same infection that lasted at least 173 days, from 08/01/2021 to 29/06/2021 (Figure 1b). She passed away on 14/08/2021 from a non-COVID related complication.

**Figure 1.**
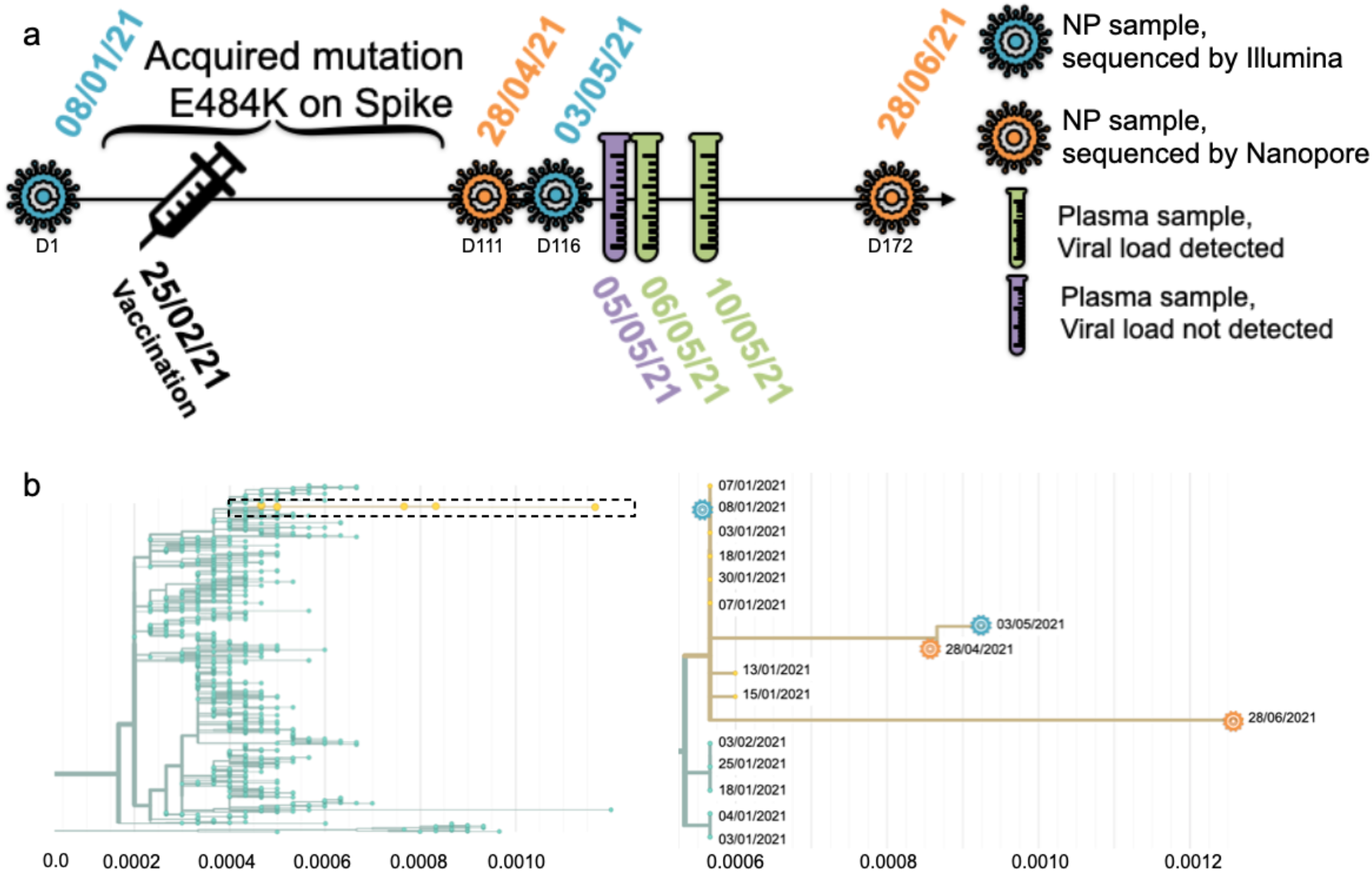
Description of Q1’s infection. **a**: Timeline of the infection and type of sample per date. NP stands for nasopharyngeal. **b**: Distance tree of lineage B.1.160 in Quebec on the left, close up of the box containing Q1’s four consensus sequences on the right. The X axis is measured in substitutions per site per year.

One immunosuppressed South Korean 25 years old male patient (K1, described as P2 in Lee et al.) was infected with PANGO lineage B.1.497 in late 2020 and early 2021.^12^ He had acute myelogenous leukemia and had received an allogeneic hematopoietic stem cell transplant one year prior. His infection lasted 73 days and 16 samples were collected over the span of the first 67 days (20/11/2020 to 26/01/2021). Neither of the two patients mentioned here were treated with mAbs or convalescent sera.^12^ The complete list of samples, dates, and tissues can be found in Table S1.

### Intra-host analysis of Q1’s samples

At D1, Q1 had all the characteristic mutations of B.1.160 in Quebec, as well as eight additional mutations (Figure 2) shared with five other sequences in the LSPQ database (Figure 1b). Sequences at D111 and D116 share ten new mutations that are not seen at D172. The additional mutations seen on D111 on Figure 1b but not in Figure 2 are low quality base calls that were filtered out from the Nextstrain analysis. The reversal of all consensus mutations acquired at D111 and D116 makes it unlikely that the substrain at D172 has evolved from the ones at D111/D116.

**Figure 2.**
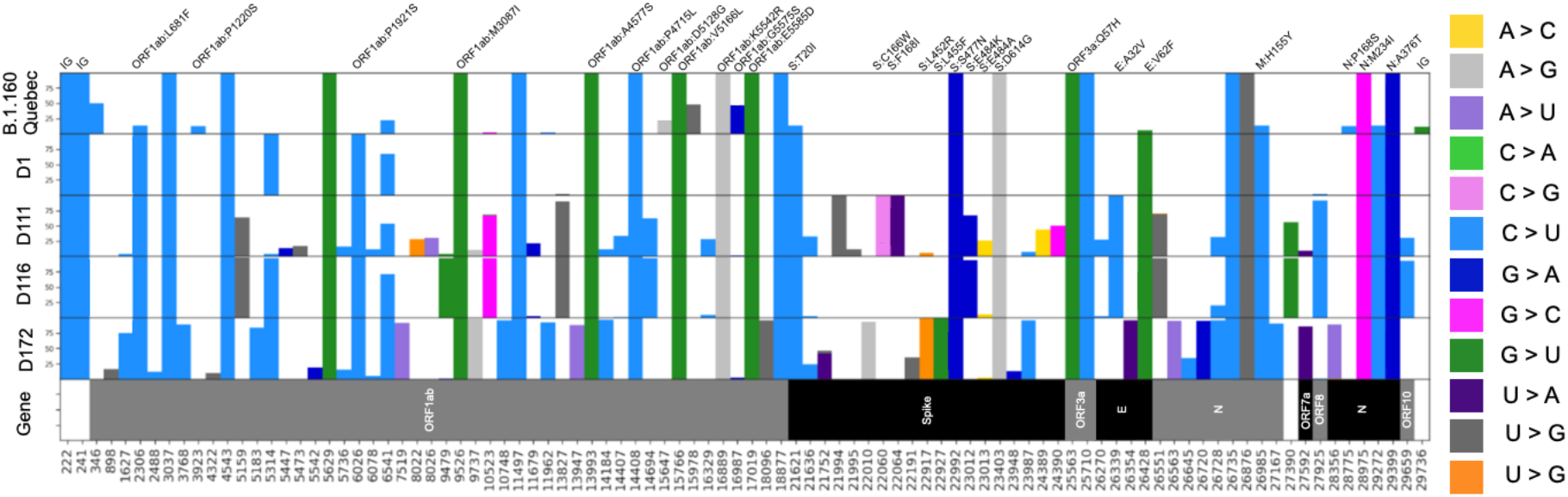
Allelic frequencies in B.1.160 in Quebec and in Q1’s infection. The top row displays the frequencies for n = 2,627 B.1.160 consensus sequences from the LSPQ database. The four following rows show intra-host frequencies for Q1’s mutations for each time point. Only mutations with intra-host frequencies above 5% for Illumina sequences (D1 and D116) and 10% for Nanopore sequences (D111 and D172) for at least one time point are presented. Because of the respective error rates of both sequencing technologies, discrepancies up to 5% for Illumina sequences (D1 and D116) and 10% for Nanopore sequences (D111 and D172) are likely to be sequencing artifacts. Non-synonymous mutations are written on top, and the color represents the nucleotide change.

### Intra-host evidence of multiple viral populations with distinct immune escape mutations

Q1 and K1 had a combined total of nine substitutions resulting in a change of five residues in the spike’s RBD (Figure 3a). Mutations at residues S:346, S:484, S:490, and S:494 confer resistance to an array of mAbs,^13^ and mutations at residue S:346 and S:348 have been linked with higher transmissibility.^14^ Both patients had substitutions G23012A and A23013C on S:484.

**Figure 3.**
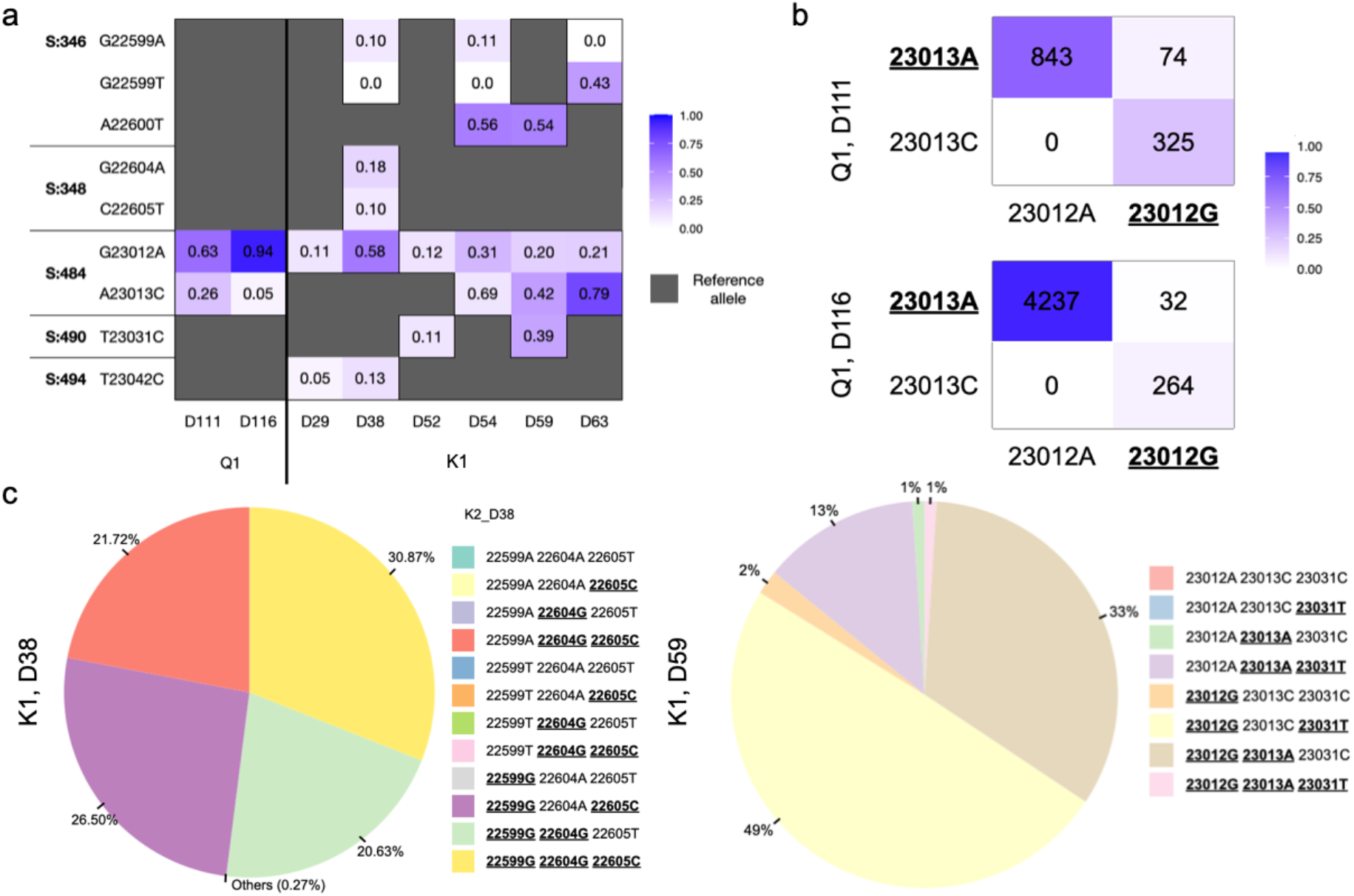
Intra-host allelic frequencies for mutated positions in the S’s RBD. **a**: frequencies for the alternative allele per position for mutated positions at S:346, S:348, S:484, S:490, and S:494 in Q1 and K1. **b**: frequencies of the haplotypes present at codon S:484 on D111 and D116 for Q1. Reference alleles are underlined. **c**: Left - frequencies of the haplotypes present at codons S:346 and S:348 on D38 for K1 on reads encompassing all three positions. “Others” category includes 22599A/22604G/22605T, 22599G/22604A/22605T, and 22599T/22604G/22605C combinations (0.1%, 0.07%, 0.1%, respectively). Right - frequencies for the haplotypes present at codons S:484 and S:490 on D59 for K1 on reads containing all three positions.

To determine the full extent of intra-host genetic diversity at a given time point, we analyzed individual reads to determine if the substitutions in the RBD belonged to different viral populations or if they co-occurred. For Q1, the substitutions on S:484 at D111 and D116 were mutually exclusive; no reads contained both alternative alleles on 23012 and 23013 (Figure 3b). There were two major distinct mutant populations of S:E484K (0.68 on D111, 0.93 on D116) and S:E484A (0.26 on D111, 0.06 on D116), as well as a small wild-type population (0.06 on D111, 0.01 on D116). For K1, G22599A and C22605T on D38 were both at a frequency of 0.10 (Figure 3a), which could suggest co-occurrence of these mutations. However, when retrieving reads containing all three positions, we see that those substitutions belong to different viral populations (Figure 3c). These results highlight the importance of analyzing aligned reads to describe the intra-host population dynamics.

### Intra-host patterns at S:E484 in the general population infected by SARS-CoV-2

Analysis of iSNVs in 147,537 SARS-CoV-2 sequencing libraries downloaded from NCBI revealed that no sequence had more than one mutation on codon S:346. Only four samples had more than one mutation on S:484 that led to distinct viral populations (SRR15258550, SRR15061404, SRR17006835, SRR16298333). No clinical details on these infections are available, but the overall mutational burden was not characteristic of protracted infections. The small number of occurrences of the RBD mutational pattern found in the general population (at a frequency below 0.003%) highlights the peculiar character of the mutational events identified in the immunosuppressed individuals described here.

## Discussion

We performed intra-host analysis on serial SARS-CoV-2 sequences from two patients with hematologic cancers and compared the identified patterns with 147,537 sequencing libraries to look for intra-host populations of immune escaping mutations in the spike’s RBD. We found evidence of multiple viral populations co-existing in this region within a single host in immunosuppressed patients. Furthermore, we found distinct populations of mutants for codon S:484 which is extremely rare in the general population, thus stressing the importance of studying immunosuppressed patients in a longitudinal design to get insights into key steps of viral evolution.

When comparing consensus mutations found at Q1’s D1, D111/D116, and D172, we saw that all four time points share the mutations present at D1, demonstrating that this is a single long-lasting infection. However, D111/D116 and D172 accumulated 10 and 20 new mutations, respectively, that are not shared across time points. The only notable exception was C26728T, a synonymous mutation in the N protein present at frequencies of 0.29, 0.2, and 0.92 on D111, D116, and D172, respectively. This suggests that the substrains present at these time points evolved separately from the substrain at D1, and that C26728T may be a recurrent mutation. The substrain present at D111/D116 was cleared from the nasopharyngeal tissues and replaced by the one at D172. Given the lack of overlap of mutations between D111/D116 and D172, the substrain present at the last time point may have evolved in another location within the host’s body, consistent with patterns observed in viral reservoirs. The theory of viral reservoirs as an explanation for long-COVID symptoms has been put forward because viral antigens or intermediate molecules of viral replication have been detected in long-COVID cases despite negative PCR tests.^17,18,19,20^ However, to our knowledge, this is the first evidence of a viral reservoir based on intra-host phylogenetics. These results call for further investigation to determine whether SARS-CoV-2 viral reservoir can be found in immunocompetent patients.

Immune escape has been a growing concern in the past year due to the rise of Omicron and its multiple sublineages that escape natural immunity as well as available vaccines and several mAbs.^19,20^ Here we described two immunosuppressed patients that were not treated with mAbs but that developed de novo multiple viral populations with mutations known to cause resistance to different mAbs.^3,6,7^ We have shown that this pattern is extremely rare in the general population, making it very likely that those distinct populations in Q1 and K1 arose due to their condition, revealing SARS-CoV-2 escape strategies. This conclusion is supported by another recent case study with an immunocompromised patient, which also found multiple mutations on S:484.^11^ As is the case for other viruses, our results suggest that combination strategies accelerating viral clearance may be required to clear viral populations with pre-existing mutations within vulnerable patients.

## Methods

### Viral databases

SARS-CoV-2 consensus sequences data were obtained from the LSPQ database through the CoVSeQ consortium (https://covseq.ca/data-info?lang=en) on 16/11/2021. Only sequences that were covered at more than 90% and a mean depth of 50X for Illumina and 16X for Oxford Nanopore technologies (ONT) with no previously documented frameshift, less than 5% N at most 5 ambiguous bases were used. Serial sequences from two patients described by Lee et al.^12^ (P1 and P2) were obtained from NCBI’s Sequence Read Archive (study SRP357108). One of them (P1), did not have iSNV in the spike’s RBD and was excluded from this study. A total of 147,537 representative SARS-CoV-2 Illumina libraries from 2020 and 2021 were downloaded from NCBI and served as a reference dataset to compare patient data (see *Intra-host analysis* below). Metadata for Q1 were obtained as part of BQC-19, PMID: 34010280.

### Whole-genome sequencing and consensus sequence generation

All LSPQ sequencing data were analyzed using the GenPipes^21^ Covseq pipelines to produce variant calls and consensus sequences. Samples were sequenced on Illumina or ONT. Regardless of the sequencing technology, data was initially processed to remove any host sequences by aligning to a hybrid reference with both human (GRCh38) and SARS-CoV-2 (MN908947.3). Any sequences that aligned to the human portion of the hybrid reference were removed from downstream analysis. For Illumina sequencing data, raw reads were first trimmed using cutadapt (v2.10), then aligned to the reference using bwa-mem (v0.7.17). Aligned reads were filtered using sambamba (v0.7.0) to remove paired reads with an insert size outside the 60-300bp range, unmapped reads, and all secondary alignments. Then, any remaining ARTIC primers (v3) were trimmed with iVar (v.1.3.4). To create a consensus representative of the most abundant species in the sample, a pileup was produced using Samtools (v1.9) which was used as an input for FreeBayes (v1.2.2). For ONT sequencing data, raw signals were basecalled using guppy (v3.4.4) with the High-Accuracy Model (dna_r9.4.1_450bps_hac). Reads outside the expected size range (400-700bp) were removed from the analysis. Reads were then aligned to the reference using minimap2 (v.2.17) and filtered to remove incorrect primer pairs and randomly downsampled to keep 800X depth per strand in high coverage regions. Finally, Nanopolish (v0.13.1) was used to call mutations in regions with a minimum depth of 16X (8X per strand) and a flank of 10bp. After masking regions with coverage below 20X, mutations called by nanopolish were integrated into the reference using bcftools (v1.9) to create a consensus sequence. In all cases, MN908947.3 was used as a reference genome. A full description of both pipelines can be found in the following URLs: https://genpipes.readthedocs.io/en/genpipes-v4.1.2/user_guide/pipelines/gp_covseq.html and https://genpipes.readthedocs.io/en/genpipes-v4.1.2/user_guide/pipelines/gp_nanopore_covseq.html.

### Phylogenetic analysis and mutational spectrum

The ‘Pangolin’ network was used to identify the sequence lineage for consensus sequences from Quebec (PangoLearn version 2021-11-09, Pangolin version 1.2.93),^22^ and all sequences characterized as the B.1.160 lineage were used to generate a distance tree. The phylogenetic trees were generated with Nextstrain viewer^23^ using the default settings.

### Intra-host analysis

The dataset for the intra-host analysis consists of sequences from one patient from the LSPQ and one patient described by Lee et al.^12^ The intra-host mutational patterns were compared to our in-house intra-host mutation database based on 147,537 representative samples. Each library was trimmed using TrimGalore! v0.6.0 and then mapped to the reference genome NC_045512.2 using bwa-mem v.0.7.17. The remaining amplicon sequences were trimmed using iVar with a hybrid amplicon definition file combining ARTIC v3, v4 and v4.1 designs. Primary reads were kept using Samtools v.1.15.1. iSNVs below 5% for Illumina and 10% for Nanopore that are not found at a higher frequency in at least one time point per patient are likely to be sequencing errors and were filtered out. Reads containing reference and alternative alleles for positions in the spike’s RBD were extracted from the BAM files using ctDNAtools.^24^ The number of reads containing different combinations of alternative and reference alleles was then compiled to determine the frequencies of the possible haplotypes.

## Acknowledgements

We thank members of the Hussin group for helpful discussions and paper revisions, specifically Matthew Scicluna, as well as Simon Gravel and Vincent-Philippe Lavallée for critically reading the manuscript. We thank Floriane Point for complementary laboratory work, and members of the Quebec public health surveillance committee for SARS-CoV-2, specifically Judith Fafard, and Canadian COVID Genomics Network (CanCOGeN), specifically Eric Fournier and Paul Stretenowich. This work was completed thanks to computational resources provided by Calcul Quebec clusters Narval and Beluga. We acknowledge and thank GISAID and NCBI as well as all contributing laboratories for giving access to their SARS-CoV-2 genome sequences. This study was supported by funding from the Canadian Foundation for Innovation, IVADO COVID19 Rapid Response grant (CVD19-030), the National Sciences and Engineering Research Council (NSERC) (ALLRP 554923 – 20), and the Canadian Institutes of Health Research (CIHR) (#174924). D.F. is a BioTalent awardee. M.C. is a FRQS Junior 1 research scholar, J.G.H. is FRQS Junior 2 research scholar and D.E.K. is a FRQS Merit research scholar. This study was also supported by the CIHR operating grant to the Coronavirus Variants Rapid Response Network (CoVaRR-Net) and the Biobanque québécoise de la COVID-19 (BQC-19). Finally, we would like to thank members of the Mila COVID19 Task Force for their camaraderie and valuable insight into data analysis strategies during the pandemic.

## Author contributions

DF, MC and JGH conceived the study, interpreted the results and wrote the manuscript. DF, RP, JCG, JHG performed bioinformatics analyses and drafted methods sections. FM, RP and JCG created and maintained the intra-host database. EBR and DK recruited Q1 and sampled SARS-CoV-2 material. EBR, SGL, NC and AP performed experiments on Q1 SARS-CoV-2 temporal samples. IL and SM supervised the CoVSeQ initiative which sequenced Q1’s SARS-CoV-2 samples. All authors revised and approved the final version of the manuscript.

## Supplementary Material

**Table S1.** Description of analyzed samples

|  | ID | Date | Days since symptom<br>onset | Tissue |
| --- | --- | --- | --- | --- |
| Q1 | L00409639001 | 08/01/2021 | 1 | Nasopharyngeal<br>swab |
|  | L00350036001A | 28/04/2021 | 111 | Nasopharyngeal<br>swab |
|  | L00350839 | 03/05/2021 | 116 | Nasopharyngeal<br>swab |
|  | L00363495001 | 28/06/2021 | 172 | Nasopharyngeal<br>swab |
| K1 | SRR17793984 | 20/11/2020 | 0 | Nasopharyngeal<br>swab |
|  | SRR17793983 | 02/12/2020 | 12 | Nasopharyngeal<br>swab |
|  | SRR17793982 | 02/12/2020 | 12 | Saliva |
|  | SRR17793981 | 02/12/2020 | 12 | Stool |
|  | SRR17793980 | 07/12/2020 | 17 | Throat swab |
|  | SRR17793979 | 09/12/2020 | 19 | Nasopharyngeal<br>swab |
|  | SRR17793978 | 09/12/2020 | 19 | Saliva |
|  | SRR17793977 | 09/12/2020 | 19 | Stool |
|  | SRR17793976 | 09/12/2020 | 19 | Urine |
|  | SRR17793975 | 19/12/2020 | 29 | Nasopharyngeal<br>swab |
|  | SRR17793973 | 28/12/2020 | 38 | Nasopharyngeal<br>swab |
|  | SRR17793972 | 11/01/2021 | 52 | Nasopharyngeal swab |
|  | SRR17793971 | 13/01/2021 | 54 | Saliva |
|  | SRR17793970 | 18/01/2021 | 59 | Nasopharyngeal swab |
|  | SRR17793969 | 22/01/2021 | 63 | Nasopharyngeal swab |
|  | SRR17793968 | 26/01/2021 | 67 | Nasopharyngeal swab |

